# A data-driven approach to integrative taxonomy

**DOI:** 10.64898/2026.07.23.740253

**Authors:** Kristian Peters

**Author notes:** Corresponding author: Kristian Peters.

## Abstract

Central to understanding biodiversity is the systematic classification of biological taxa where concepts on integrating ‘omics data have not yet been comprehensively executed. The data-driven approach is aiming at integrating phylogenetic data from three or more scales aiming at enriching the robustness of species delimitations while also revealing evolutionary drivers of diversification and speciation.

This study investigates complex-thallose liverworts as reference and combines the analysis of DNA marker sequencing, morphometrics utilizing bioimaging measurements and chemometrics using liquid chromatography high-resolution mass-spectrometry (UPLC/ESI-QTOF-MS) with data-dependent acquisition of tandem mass-spectra (DDA-MS), with character variation evaluated with dendrograms and data mining.

The comparative analysis revealed similar tree topologies albeit with diverging positions of taxa that were attributed to flavonoid-glycosides, auronidins and phenanthrenes resulting from evolutionary radiation and adaptations to stressful bioclimatic conditions, whereas unique fatty acyls in *Riccia* link to ecological causes of speciation. Characteristic amino acid motifs confirm the intermediary position of liverworts between algae and land plants.

By functionally attributing and mapping chemometric markers to the taxonomy, the data-driven approach integrating ‘omics into integrative taxonomy is opening opportunities for plant systematics to identify evolutionary drivers and form new hypotheses on the diversification and speciation of species.

## Introduction

Plant systematics is central to understanding and characterizing biodiversity^1^. Taxonomy aims at systematically describing, classifying and naming biological taxa, including species and higher-ranking orders in the taxonomic ontology. As species are the indispensable units to assess biodiversity, in the light of the current accelerating loss of biodiversity this has led to the urgent need for more realistic taxonomic circumscriptions and insights into evolutionary mechanisms^1,2^. Since about 30 years, empirical DNA barcoding methods (marker sequencing) have revolutionized plant systematics fundamentally reshaping taxonomic inference and largely superseding classical morphology-based approaches^3^. Since then many investigations have almost exclusively focused on DNA barcoding methods for reconstructing phylogenetic relationships^3^ leading to limited species circumscriptions often disregarding character evolution at other than genetic scales such as biochemical, morphological, or ecosystem scales^4^. This has led to the development of integrative taxonomy that aims at combining different often domain-agnostic data^5^ and necessitates to delimitate species based on hypotheses from several domains and requires interdisciplinary knowledge. As evolution of taxonomically relevant characters is often challenged by simultaneous mechanisms (such as speciation rate, character variation, different environmental and biotic selection processes, biological organization) acting differently at different scales, this has become a challenging task^1^.

With the rise of affordable NGS and ‘omics methods, the field is about to be revolutionized again. However, the wealth of information provided by ‘omics allows conclusions on evolutionary mechanisms that go well beyond pure plant systematics. In recent years, interest in morphometrics in explaining species relations is rising again due to also capturing ecological factors such as environmental changes and due to the emergence of AI to greatly facilitate analyses of characters in images^6^. In addition, chemometrics as a tool for plant systematics has been long underestimated, especially with development of high-throughput metabolomics annotation techniques^7^. In plant systematics, this plethora of information at different scales has not yet been fully integrated, leaving opportunities to resolve the ecological circumstances that have led to evolutionary processes such as diversification or extinction.

The data-driven approach to integrative taxonomy combines classical morphometric and marker sequencing with ‘omics and enables not just insights into (1) evolutionary relationships among taxa to produce more robust phylogenies, but also (2) the evolutionary drivers underlying processes such as diversification or speciation. Along DNA barcoding sequences, additional phenotypic data in the form of bioimages are available that can be formulated as taxonomic characters by performing morphometric measurements such as determining plant stature length, leaf characteristics, or cell lengths coupled with statistical analyses. In addition, ‘omics techniques such as eco-metabolomics^8^ captures information that need to be attributed to evolutionary or ecological mechanisms revealing new mechanistic insights that go beyond purely taxonomical discrimination. However, data should not be integrated blindly and representative characters chosen that largely reflect phylogenetic properties at the respective scale. Furthermore, characters need to be converted to compatible metrics from which phylogenetic trees can be calculated.

The data-driven approach is aiming at integrating phylogenetic data from three or more scales aiming at enriching the robustness of phylogenetic consensus trees when sequence-based data is scarce or incomplete, resolving taxonomic or phylogenetic disparities or circumscribing ecotypes when phenotypic plasticity is large. In addition, it enables insights into evolutionary mechanisms and the generation of new hypotheses that can be tested in follow-up experiments.

To demonstrate the data-driven approach to integrative taxonomy, a reference dataset of bryophytes (complex-thallose liverworts, order Marchantiales) is reused. The methods described herein are also applicable to other kinds of organisms. In contrast to cryptogams such as lichens which traditionally integrate chemometrics and marker sequences^4^, with bryophytes there has been a strong emphasis on DNA marker-based investigations despite that traditional Sanger-based sequencing of core-genomes in certain groups such as liverworts are very challenging^7^. Although shotgun whole-genome sequencing like Illumina, Nanopore or Nanoball became affordable in the past years, there is still hesitancy to use these new and more complete methods for evolutionary investigations probably due to missing reference genomes, complex bioinformatic processing and due to the fact that there are only few economically relevant bryophytes. Only in recent years, the comparative study of morphological characters has gained attention again for bryophytes^9,10^. Still, morphological characters are often treated separately rather than fully integrated and used merely to confirm or disregard evolution at the genetic scale. This is mainly due to the fact that recent scientific developments have rarely been brought forward to bryophyte systematics. By contrast, for vascular plants there have been tremendous progress in bioimage analyses that allow for the semi-automated assessment and morphometrics of characters^6^. In contrast to lichens where chemometrics have been applied for over 50 years, reconstructions of phylogenies based on chemometric markers were rarely accomplished with bryophytes^11–14^. Given the challenges with traditional methods it has become a viable alternative^8^. Chemotaxonomy that investigates chemometrics to reconstruct phylogenetic relationships is based on the assumption that species have a constant core metabolome, irrespective of their geographic origin, or ecology^8^. In the past 10 years, there has been great progress in assessing and annotating chemometric markers using untargeted metabolomics^15^. The analytical instrumentation and computational methods in metabolomics have emerged to be nearly as complete as shotgun genomics methods and are comparably affordable in assessing phylogenetically relevant markers at the biochemical scale^8^.

This study investigates complex-thallose liverwort species of the order Marchantiales in biological soil crusts, which are of particular interest in studying early land plant evolution. They are among the oldest living relatives of early land plants and evolved in the Devon approx. 450 Ma ago^16^. In their dominant gametophytic stage, they possess key innovations of land plant evolution, including a shoot apical meristem yielding a 3-dimensional thallus with specialized types of cells providing the key morphological and physiological traits for terrestrial adaptation^17^. Furthermore, flavonoid biosynthesis, key biochemical constituents for land adaptation, have also been evolved and radiated in this group^18^. The late Permo-Triassic (approx. 200 Ma ago) gave rise to the Ricciaceae^19,20^ which coincided with massive ecological reorganizations and extreme conditions as part of the Triassic-Jurassic mass extinction event. These rather harsh conditions are still selection factors in biocrusts today^21,22^. As a result, the investigated species have developed adaptive traits and life-history strategies to withstand desiccation, temperature extremes and high levels of radiation^23^.

16 samples of 13 species were investigated from biological soil crusts in Southern Sweden and Central Germany. Whereas nine of the investigated species belonged to the family of Ricciaceae, the outgroup was represented by four species belonging to Aytoniaceae and one to Cleveaceae within the order of Marchantiales (Fig. 1, Table 1). *Riccia subbifurca* has been sampled twice (Fig. 1o,p) and *R. ciliifera* (Fig. 1i,j) in Sweden and Germany (Fig. 1), respectively. The data-driven approach to integrative taxonomy is based on the assessment of taxonomically-relevant characters at three distinct scales: DNA barcoding (*trn*LF markers), morphometric cell and thallus measurements using deterministic bioimage analyses and chemometrics using untargeted liquid chromatography high-resolution mass-spectrometry (UPLC/ESI-QTOF-MS) coupled with data-dependent acquisition (DDA-MS)^8^. Using statistics and machine learning, morphometric and chemometric characters that have high phylogenetic relevance were extracted. To contextualize the information at different scales, (1) morphometric, (2) chemometric and (3) sequence-based traits were integrated with each other. It is hypothesized that chemometric characters capture more phylogenetically relevant information than morphometric characters and that the morphometric and chemical information are vital additions to sequence-based markers and enable to better understand species delimitation and causes of diversification^14,24–26^.

**Figure 1.**
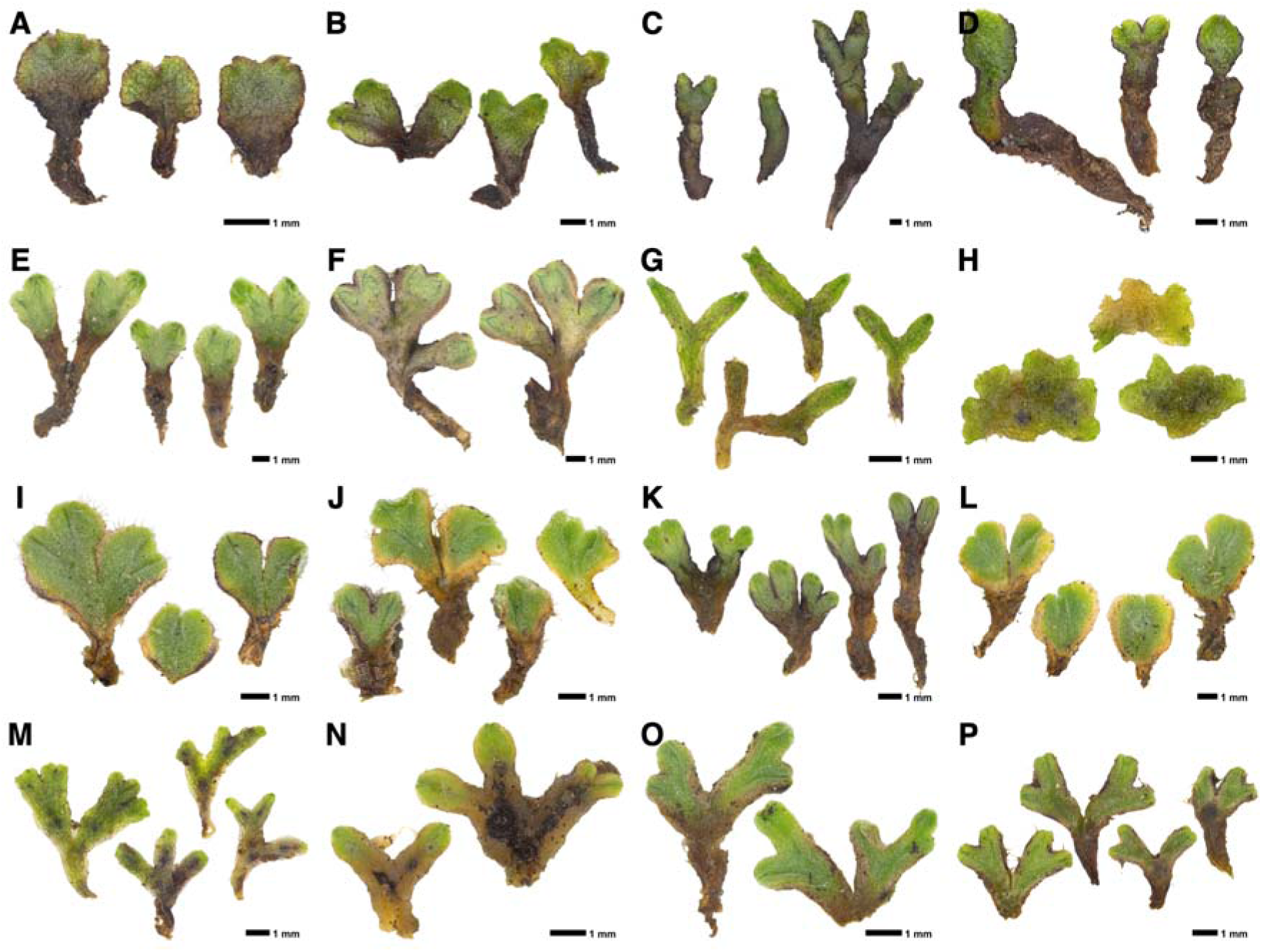
Stature (ventral sides) images and overview on the investigated specimens. **(a)** *Asterella gracilis* (F.Weber) Underw. **(b)** *Athalamia hyalina* var. *suecica* (Lindb. ex Gottsche & Rabenh.) S.Hatt. **(c)** *Mannia fragrans* (Balb.) Frye & L.Clark **(d)** *Reboulia hemisphaerica* subsp. *hemisphaerica* (L.) Raddi **(e)** *Riccia beyrichiana* Hampe **(f)** *Riccia bifurca* Hoffm. **(g)** *Riccia canaliculata* Hoffm. **(h)** *Riccia cavernosa* Hoffm. **(i)** *Riccia ciliifera* Link GER **(j)** *Riccia ciliifera* Link SWE **(k)** *Riccia gothica* Damsh. & Hallingb. **(l)** *Riccia ciliifera* var. *gougetiana* GER **(m)** *Riccia huebeneriana* Lindenb. **(n)** *Riccia sorocarpa* Bisch. **(o)** *Riccia subbifurca* Warnst. ex Croz. SWE1 **(p)** *Riccia subbifurca* Warnst. ex Croz. SWE2.

## Results

### Phylogenetic trees at their respective scales

Phylogenetic trees were constructed from the marker sequences (Fig. 2a), chemometric (Fig. 2b) and morphometric characters (Fig. 2c). The trees were calculated *de novo* with RAxML-NG using the model GTR with discrete GAMMA (GTR+G). Alpha of phylogenetic trees was estimated using Maximum Likelihood (100) and clade support were estimated using bootstrapping (1000). Tree topologies were slightly different between the scales whereas the genetic scale produced the most and the morphometric scale the least realistic representation of taxonomical relationships (Fig. 2). The phylogenetic relationships in the trees constructed at the different scales were largely conserved regarding the outgroup (non-Riccia liverworts *At. hyalina*, *As. gracilis*, *M. fragrans*, *Re. hemisphaerica*), Ricciella (*R. canaliculata*, *R. huebeneriana*, *R. cavernosa*), and Eu-Riccia (*R. ciliifera*, *R. sorocarpa*, *R. bifurca*, *R. beyrichiana*, *R. subbifurca*, *R. gothica*) groups albeit with different positions and confidence (Fig. 2a-c). The positions of *R. subbifurca*, *R. gothica* and *R. canaliculata* were intermixed in the tree trees leading to inconclusive phylogenetic positions at the different scales.

**Figure 2.**
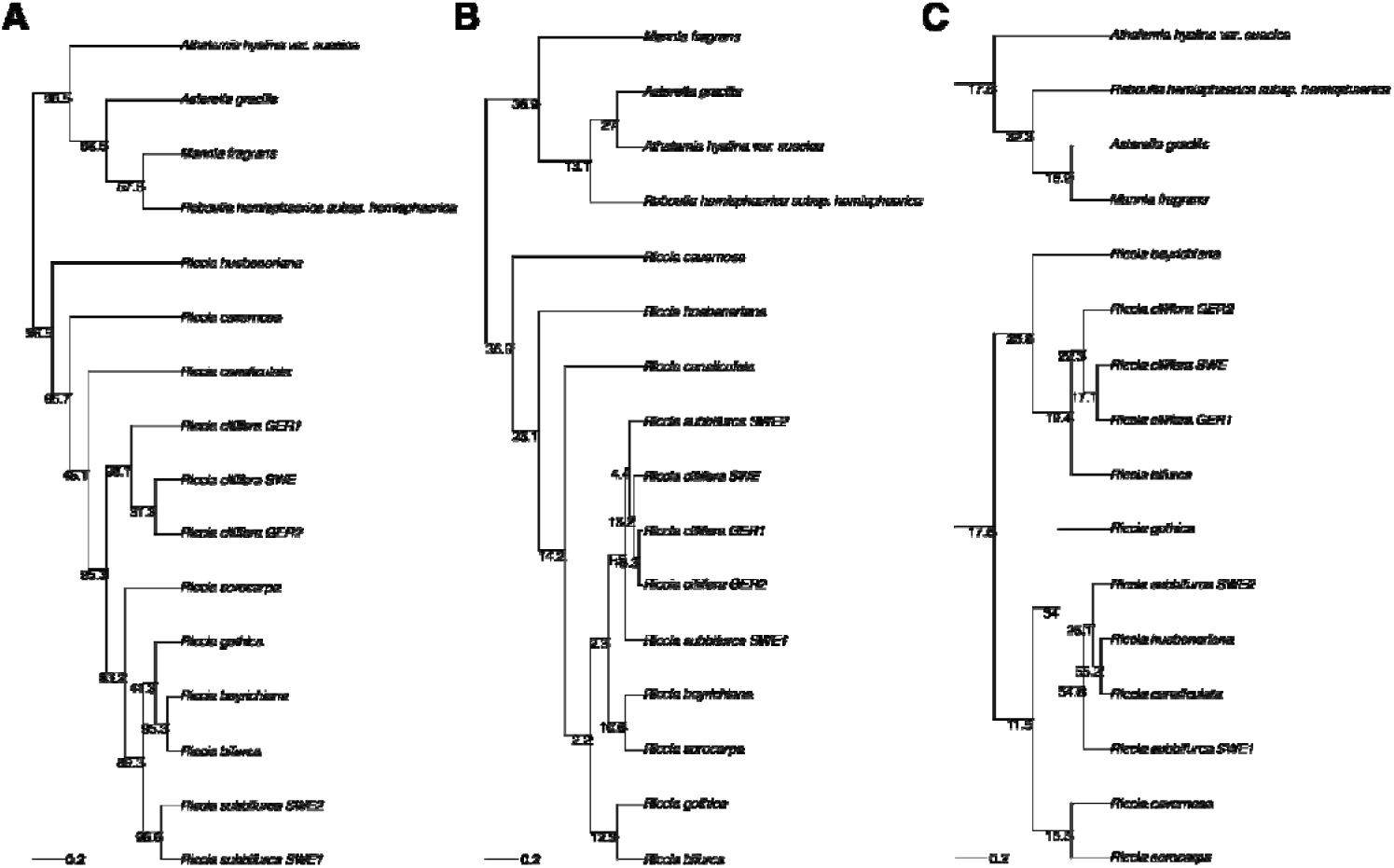
Individual phylogenetic trees constructed from the **(a)** *trn*LF marker sequences, **(b)** chemometric, and **(c)** morphometric measurements. The numbers on the branches indicate branch confidence. The edge length is given in the bottom left corner.

### Integrative taxonomy improves species delimitation

Next, a comparative plot of the three phylogenetic trees in Figure 2 was accomplished (Fig. 3). Manually inspecting the differences in the trees revealed the marker-based and chemometric trees (RF=0.643) to be more similar than to the morphometric tree (RF=0.786) (Fig. 3). The largest displacements were found for both *R. subbifurca* specimens in the chemometric tree and for *R. canaliculata* in the morphometric tree indicating adaptations to different abiotic or biotic conditions rather than phylogenetic differences.

**Figure 3.**
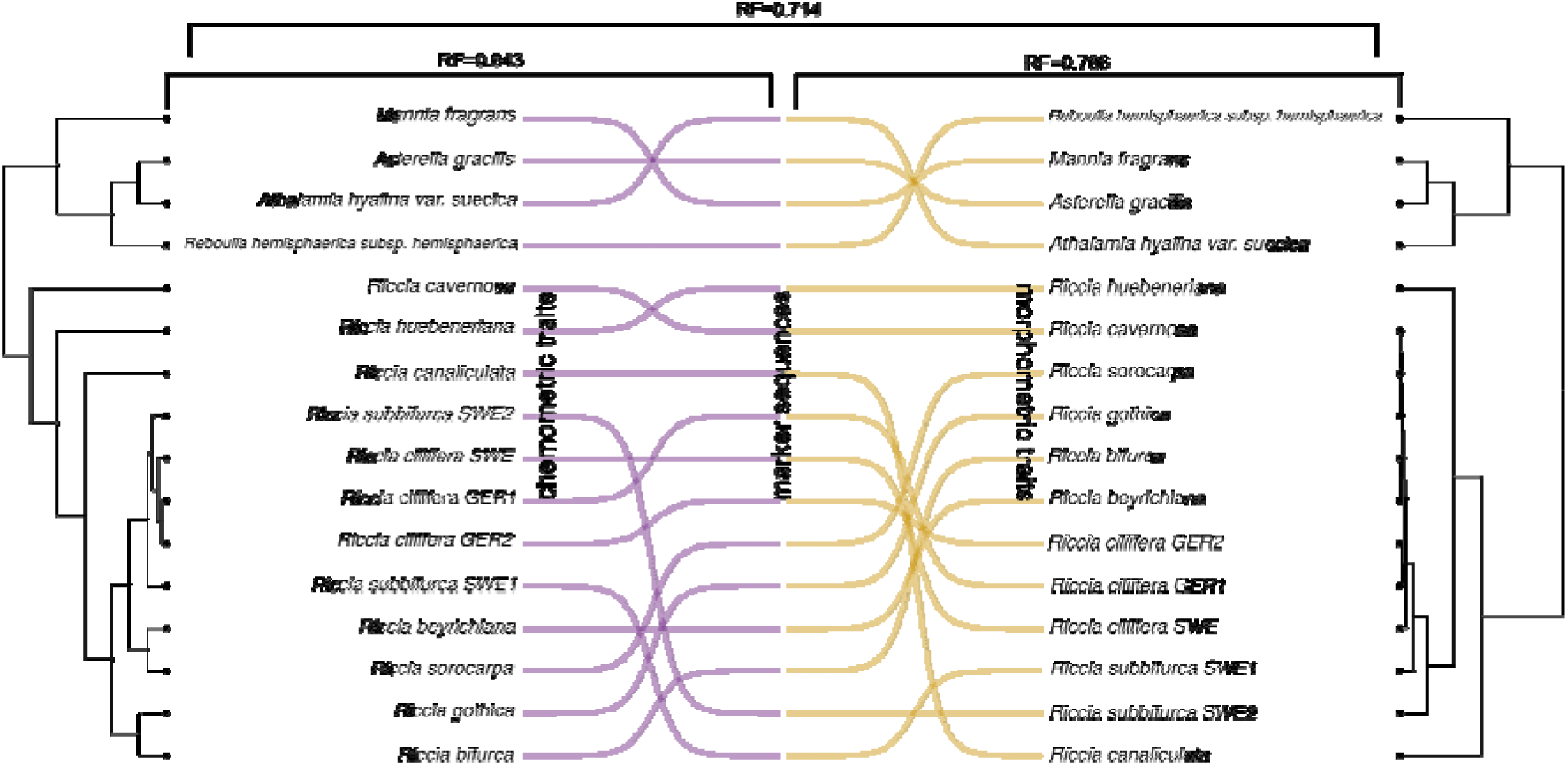
Comparison of trees obtained from the selected molecular traits (at the level of compounds) (left, purple color), the *trn*LF sequences (center), and the measured phenotypic traits (right, dark yellow color). Species names and dendrogram resulting from markers are not shown in the center. Robinson-Foulds similarity of molecular traits and sequence trees: RF=0.643. Robinson-Foulds similarity of phenotypic traits and sequence trees: RF=0.786. Robinson-Foulds similarity of molecular and phenotypic traits: RF=0.714. The lower the value the more similar the trees.

As the information based solely on plastid *trn*LF marker sequences did not produce a tree confident enough to delimitate the different species (Fig. 2a), the chemometric and morphometric information were integrated with sequences to improve information and distances between species profiles. This was accomplished by constructing a *de novo* consensus tree without prior knowledge of species delimitations (Fig. 4). The consensus tree was constructed using IQTREE and the MAST algorithm calculating 1000 bootstrapped trees obtained from the respective scales. To not bias the construction of the consensus tree and to have a similar number of character states at the respective scales, only the 208 most significant chemometric characters selected by the Random Forest model (Fig. 7 below) were chosen. The consensus tree (Fig. 4) showed an improved and more realistic representation of species delimitations^27^ as taxa positions, distances and edge lengths to and within the outgroup were enlarged when compared to the sequence-based tree (Fig. 2a). The three specimens of *R. ciliifera* and the two specimens of *R. subbifurca* were clustered closely together confirming their phylogenetic identity. The *Riciella* group and *R. cavernosa* were closer to the outgroup confirming prior phylogenetic investigations.

**Figure 4.**
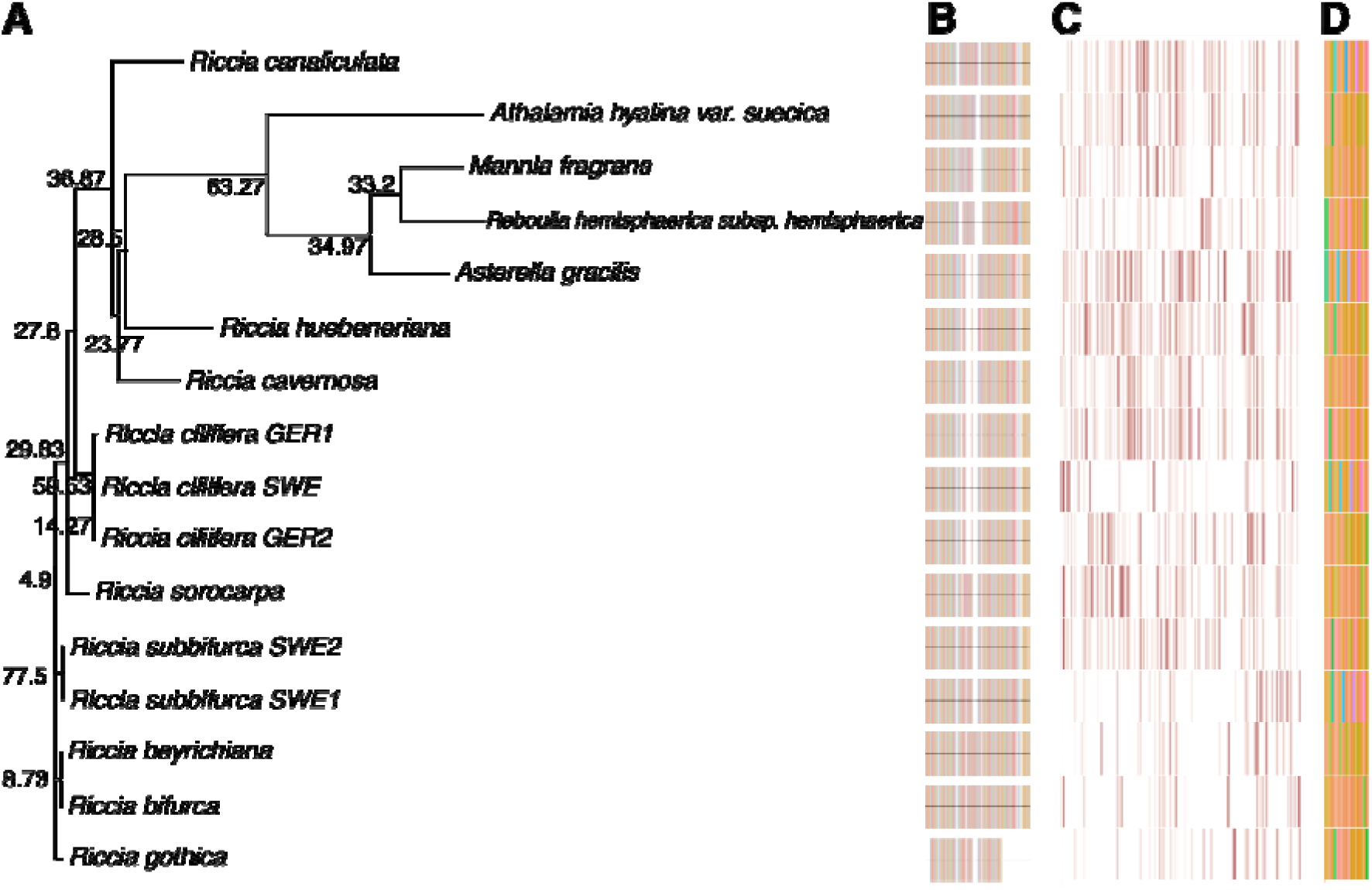
**(a)** Consensus tree on the left side constructed from the sequence-based, chemometric and morphometric data. The numbers on the branches indicate branch confidence. The right side shows the **(b)** marker-sequence data, **(c)** 208 selected chemometric markers, **(d)** morphometric markers.

### Scale-dependent phylogenetic divergence

The rates and similarities of the phylogenetic divergences in the species were assessed with PCA plots (Fig. 5). Divergences were different at the respective scales, whereas distances are most pronounced at the sequence-based scale (43.88% explained variation on PC1+2) and least at the morphological scale (14.84% explained variance) (Fig. 5a,c). The chemometric scale retained an intermediary position (37.84% explained variance) (Fig 5b).

**Figure 5.**
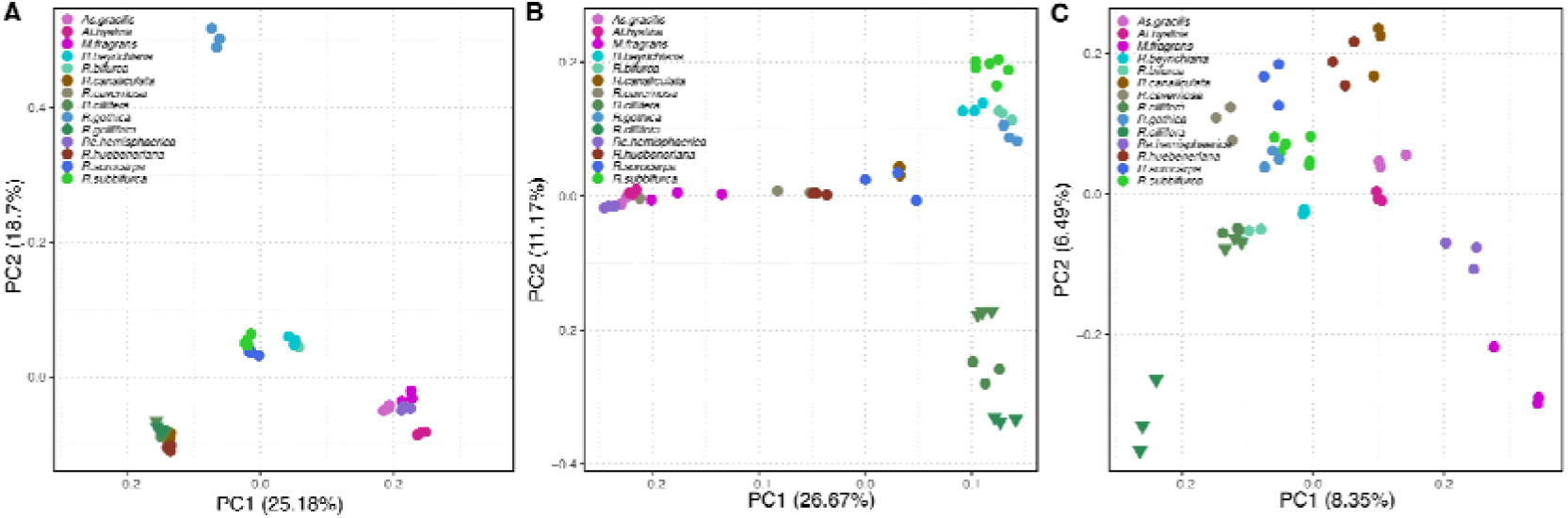
Principal Component Analysis (PCA) of **(a)** marker sequences, **(b)** chemometric, and **(c)** morphometric measurements. The x and y axes show the first two principal components. The amount of explained variation on the corresponding axes is given in brackets. Taxa have been colored.

### Scale-dependent taxonomic relevance

To quantify the variability explained by either phylogenic or ecological factors, variance partitioning was employed on the matrices with taxonomic identities and ecological factors extracted from the BET dataset^28^ (Fig. 6). At the sequence-based scale, most variance is explained by species, whereas residuals are largest at the morphological scale (Fig. 6a,c). Temperature in the coldest months, mean annual air temperature and growing degree days were the most important ecological factors defining evolutionary disparity between the investigated taxa. Life strategy and life history were only marginally important. It should be noted that evolutionary processes could not be fully separated from ecological processes and, thus, part of variance was likely shared among investigated factors.

**Figure 6.**
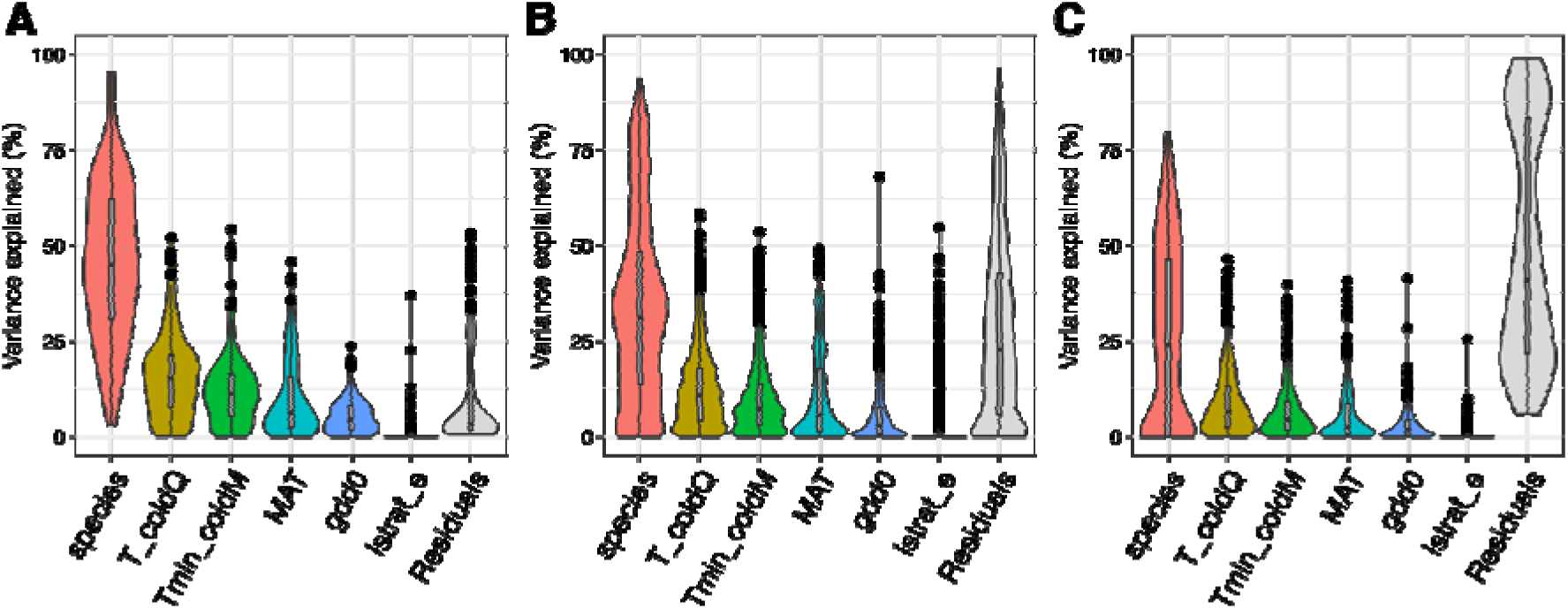
Variance partitioning. Violin plots showing the variance partitioning and the explained variance of the species and the top-five selected ecological factors for the **(a)** marker sequences, **(b)** chemometric, and **(c)** morphometric measurements. The y axis shows the amount of variation explained. The x axis shows 5 ecological factors that explain the most variation: T_coldQ: Temperature in the coldest quarter, Tmin_coldM: mean daily minimum air temperature of the coldest month, MAT: mean annual air temperature, gdd0: growing degree days heat sum above 0°C, lstrat_e: During’s life strategy, extended system, Residuals: unexplained variation.

### Integrative taxonomy reveals evolutionary drivers

To gain deeper insights into the patterns and context dependencies of chemometric markers, a random forest (RF) model discriminated 208 molecules in the investigated species (R^2^=1.0, multi-class area under the curve=0.91) (Fig. 7a). Putative annotation and chemical classification of selected molecules were performed with SIRIUS and CANOPUS^29–31^. Molecules were attributed to molecular and ecological function using the approach described in Peters et al. (2026)^32^. Molecules were then generalized into four functional categories: (1) Metabolic compounds that were involved in cell metabolism and homeostasis within plants, (2) Biological activity characterizing compounds to have biotic relationships, (3) Compounds involved in biosynthesis and plant growth, and (4) Adaptation to abiotic conditions, e.g. changing environmental conditions. To honor the multi-functionality concept^33^, association of molecules to several categories was possible. The selected molecules were set in context to the phylogenetic trees at the respective scales (Fig. 2a-c,3,4) to reveal their phylogenetic evidence.

**Figure 7.**
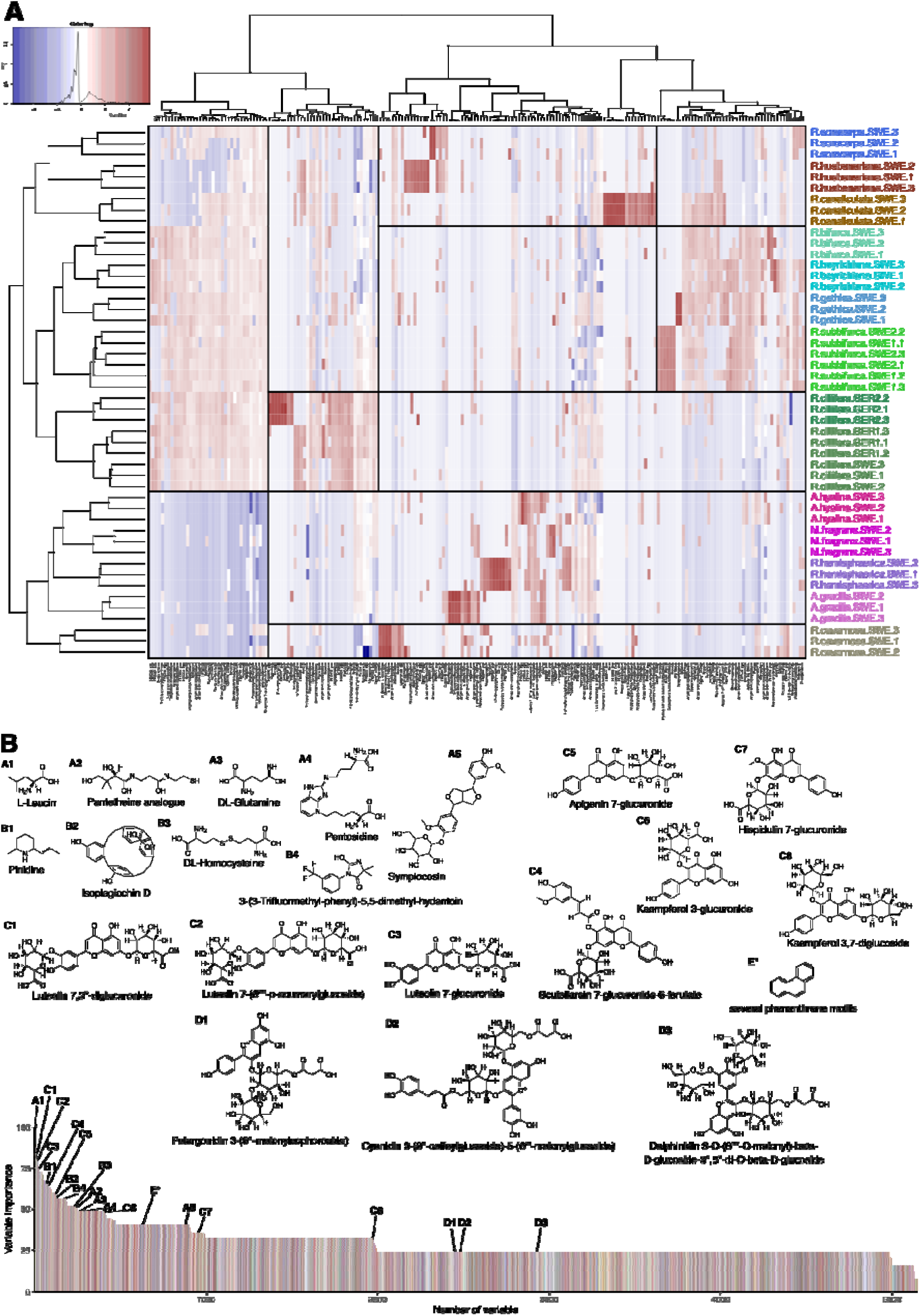
**(a)** Heatmap showing the abundance and distribution of 208 by RF selected chemometric characters in the different species. On the x axis the molecules are shown. The y axis contains the samples colored for species. Clusters on both axes were generated using Euclidean distance measure and the Ward clustering method. A red color indicates up-regulation and a blue color down-regulation. A white color means that the molecular feature was not detected. **(b)** Variable importance of molecular features harmonized to percentage values sorted by their importance with the most important on the left. The structures and the importance of some remarkable molecules are shown.

43.7% of the selected molecules were found to be involved predominantly in intrinsic cellular regulation processes uniquely characterizing the metabolism of the species and keeping cellular homeostasis comprising various alkaloid derivatives, steroids, prenol lipids, benzenes, amino acids and peptides (Fig. 7). Among the annotated molecules with highest variable importance were the amino acids L-Leucin (A1 in Fig. 7b), Pantetheine (A2), Glutamate (A3), Pentosidine (A4) and Symplocosin (A5). 33.3% of the selected molecules were annotated to have biological activity such as the only macrocyclic bisbybenzyl Isoplagiochin D (B2), Pinidine (B1), DL-Homocysteine (B3) and a Hydantoin derivative (B4). 23.0% of molecules were attributed to extrinsic bioclimatic processes linking evolutionary mechanisms to ecological processes such as environmental adaptation, plant growth, photo- and biosynthesis (Fig. 7). Here, the investigated liverworts produced an extraordinary diversity of flavonoid-O-glycosides with species-specific abundance like Luteolin 7,3’’-diglucuronide (C1), Luteolin 7-(6’’’’-p-coumarylglucoside) (C2), Luteolin 7-glucuronide (C3), Scutellarein 7-glucuronide-6-ferulate (C4), Apigenin 7-glucuronide (C5), Kaempferol 3-glucuronide (C6), Hispidulin 7-glucuronide (C7) and Kaempferol 3,7-diglucoside (C8). Bryophyte-specific phenylpropanoids including auronidin-like flavonoid pigments (D1-D3) and several molecules with complex phenanthrene-motifs (E*) were identified and had direct relationship to the presence of purple colors in *R. ciliifera*, *R. subbifurca* and *R. beyrichiana*.

Selected molecules were also investigated for their affinity to natural product classes (NPC)^34^. Species identity was largely determined by the composition and abundances of alkaloid, steroid and terpenoid classes. Fatty acids, fatty acyls and lipids were especially characteristic in *Riccia*. Non-*Riccia* liverworts were particularly rich in flavonoids, stilbenes, lignans and other bibenzyls. This affinity to natural product classes was also reflected in the structure of the phylogenetic tree (Fig. 2b).

## Discussion

The data-driven integrative taxonomy enabled the construction of more robust phylogenetic trees and improved the phylogenetic inference of taxa by having more data available to discriminate and characterize the taxa. In this study, the MAST algorithm in IQTREE was chosen for consensus tree construction^35^. Other algorithms such as Jackknife analysis^36^ or total evidence analysis^37^ may be investigated for better representation of underlying metrics at the respective scales. In addition, several metrics have been proposed to assess the level of topological congruence between two or more phylogenies such as the Kishino-Hasegawa test^38^, Likelihood Ratio (LR) test^38^, Incongruence Length Difference (ILD) test^39^, ILD test for pairwise comparisons^37^, Partition Homogeneity Test as implemented in the software PAUP*^40^ or the Templeton test^41^. Even different approaches such as taxology^42^ that are beyond the scope of this study can be explored to be used for data-driven integrative taxonomy approaches.

By investigating the different phylogenies comparatively, differences in the evolutionary significance of characters can be revealed at the respective scales as well as confirmations of the evolutionary positions of taxonomical groups. The position of the outgroup containing *At. hyalina*, *As. gracilis*, *M. fragrans* and *Re. hemisphaerica* was conserved at all three scales. The different positions of *R. canaliculata* at the periphery of the morphometric tree indicates very distinct morphological characters such as the size and structure of air pores and epidermis cells that likely represent adaptations to its semi-aquatic lifestyle when compared to the other taxa. In particular, due to the large chemometric divergence, *R. gothica* was positioned at the periphery of the consensus tree confirming recent investigations in the genus *Riccia*^27^. The disparities at the different scales show that a comprehensive choice of representative characters is necessary to produce robust phylogenetic trees^43^. As each scale also captures ecological variation that cannot be fully separated from evolutionary processes, a weighting might be performed with integration of consensus trees using intermediate penalties for multistate characters at the respective scales^44^. For example, chemotypes of lichens were found to be more consistent with phylogeny than morphological characters^45^. Here, a down- or up-weighting might produce more robust consensus trees. In summary, the consensus tree produced in this study showed a more realistic representation of the actual phylogenetic relationships^46^ as distances between and within groups of taxa as well as in edge lengths were improved. This result is in line with other observations in liverworts combining morphology and marker sequencing^20^. Metabolomics used in this study resulted in 208 chemometric characters that were significantly up- or downregulated in the investigated taxa and groups of taxa. While a descriptive analysis of the relationship of these molecules in the different taxa revealed distinctive phylogenetic relationships, a functional analysis attributing the selected molecules with either evolutionary or ecological functions revealed the underlying drivers and mechanisms underlying the taxonomical divergences. In the following, these functions are described in detail.

Overall, 43.7% of annotated molecules were functionally related to evolutionary processes. Amino acids such as leucin and the presence of leucin motifs in peptide and protein precursors were especially abundant in Ricciaceae (except *R. subbifurca*). They regulate protein synthesis, are involved in early land plant signaling, cell growth, or non-host specific plant defense^47^. Leucin may further be indicative for early plant development and proliferation because thallose liverworts like *Mannia* or *Reboulia* are usually growing in spring and summer seasons unlike the investigated *Riccia* which are annual liverworts predominantly developing in autumn. Several other related amino acid motifs were identified explaining mechanisms of early land plant evolution such as Pantetheine (A2), a pantothenic acid analogue, which is involved in the vitamin B5 biosynthesis but may also inhibit germination and growth of vascular plants^48–50^; Glutamate (A3) which has various functions in plants regulating cell homeostasis, being part of biosynthesis and catabolism pathways, involved in the uptake and storage of organic N, or as a signaling molecule regulating shoot organogenesis and stress response^51,52^; the amino acid Pentosidine (A4) involved in protein glycation in plants^53^; the glycylated triterpene saponine Symplocosin (A5) which is produced by a variety of plants as protection against pathogens^54^.

33.3% of annotated molecules were attributed to have biological activity. These molecules may indicate co-evolutionary processes between plant and associated microbial species in biocrusts but their modes of action need to be confirmed by follow-up experiments as associated microbials were not investigated in this study. Noteworthy were the macrocyclic bisbybenzyl Isoplagiochin D (B2) typically for Marchantiaceae liverworts that exerts strong cytotoxic activities, especially in fungi^55,56^. Remarkable were also molecules indicating interactions with non-microbials such as the piperidine alkaloid Pinidine (B1), previously only found in gymnosperms (Pinaceae), with anti-insecticidal properties^57^, DL-Homocysteine (B3) as part of the redox-sensitive methionine metabolism may also have a role in the response to environmental stress^58^ and a potentially nematocidal Hydantoin derivative (B4)^59^.

23.0% of molecules were attributed to relate to bioclimatic processes linking molecular mechanisms in plant growth, photo- and biosynthesis to abiotic functions such as environmental changes. Here, the investigated liverworts produced an extraordinary diversity of flavonoid-O-glycosides with species-specific abundance (C1-C8) predominantly serving as UV-B protection agents, antioxidants, or growth regulators whereas the type of glycosylation modifies their biological activity^60,61^. Biosynthesis is mediated by genes such as MpUGT735A2 and MpUGT743A1 identified in *Marchantia* yielding UDP-glycosyltransferases as part of specialized side chains of the core phenylpropanoid pathway^62,63^. These molecules are likely not phylogenetically conserved as plants produce a large diversity of different flavonoids that are flexibly glycosylated depending on the bioclimatic conditions. Bryophyte-specific phenylpropanoids including auronidin-like flavonoid pigments (D1-D3) and several molecules with complex phenanthrene-motifs (E*) were identified that likely act as a response to UV-B light stress (flavonoids) and desiccation (phenanthrenes)^18,64–67^. Indeed, D2 and D3 were detected in *R. canaliculata* and *R. ciliifera* – the only *Riccia* species with violet pigmentation in ventral scales confirming their role as pigments^66^ and investigations at the morphometric scale. In addition, the structure D1 was identified also in *R. subbifurca*, and *R. beyrichiana* that also can have facultative violet coloration in parts of the outer layer of the thallus.

Fatty acids, fatty acyls and lipids were especially characteristic in *Riccia* coherent with earlier findings^26,68,69^. The high diversity of selected molecules in the classes mycosporines, coumarins and carotenoids indicate the importance of species-specific molecular responses as well as evolutionary similarities of liverworts in the composition of classes produced by algae^70,71^. In addition, non-*Riccia* liverworts were especially rich in flavonoids, stilbenes, lignans and other bibenzyls confirming their closer phylogenetic relationship to *Marchantia*^17,72^. Contradictory to earlier investigations of liverworts, different and rich sets of alkaloids were annotated^65^. Lastly, macromolecular phenolic tannins were only found in *Riccia cavernosa*, which produces a strongly porose thallus with many air chambers which likely make the plants more susceptible to pathogenic infestation. Thus, it is hypothesized that in liverworts tannins may serve as protection against mechanical damage and infection by microbials or insects similarly to vascular plants^73^. Moreover, tannins may be a unique evolutionary invention in the cell wall of some liverworts that warrants closer investigation^65,74^.

Furthermore, species differed considerably in the composition and abundance of structurally related flavonoids, glycosides, and fatty acyls (*Riccia*) and prenol lipids (non-*Riccia*) suggesting molecular diversification due to different bioclimatic conditions^18,68,75^. This has likely giving them evolutionary advantages in the different habitats whereas molecules underlying bioclimatic traits also represent immediate and flexible molecular responses to short-term environmental changes^65^. As shown for flavonoids, molecules are likely to carry out multiple cellular functions due to different (post-)transcriptional regulations, including protein confirmations and glycosylation, as well as in ecology where molecules likely serve multiple functions which increases resilience to stress and overall fitness, but exacerbating the functional and mechanistic characterization^33,76–78^. Kaempferol shows antifungal activity and plants increase to avoid fungal infestation or boost endophytic bacterial productions^79^.

In conclusion, the data-driven approach to integrative taxonomy presented herein demonstrated that by fully integrating classical DNA plastid markers with morphometric and chemometric data improves the reconstructions of phylogenetic relationships among taxa and across groups, as well as providing mechanistic insights into ecological and evolutionary circumstances that may have led to diversification or speciation. Full integration necessitates a functional annotation of molecular characters that is following the scheme: (1) identifying significant characters, (2) a functional annotation of the selected characters revealing biological and/or ecological roles, and (3) mapping the characters to the phylogenetic tree. For example, among the molecules with largest variable importances and structural diversity were flavonoid-glycoside derivatives, auronidins and phenanthrenes that result from less-conserved modified side chains of the highly conserved core phenylpropanoid pathway representing, both, diversification resulting from past evolutionary radiation and the biochemical reservoir of liverworts to the highly stressful bioclimatic conditions in biocrusts^18,80–82^. Similarly, the characteristic production of fatty acyls (*Riccia*) and prenol lipids (non-*Riccia*) mediating similar functions and are less abundant in vascular plants and other bryophytes^65^ suggests molecular selection due to different bioclimatic conditions^18,68,75^. The large diversity in these compounds likely translates into functional divergence and may explain the ecological conditions under which the species, both, diversify and also speciate^68,83^. Moreover, a characteristic composition of amino acid motifs in precursors of peptides and proteins was found that were directly attributed to early land plant evolution validating the intermediary position of liverworts between algae and land plants^17,84^. As shown in this study, similar to the annotatable gene space^85^, the detailed information from functionally annotating chemometrics data integrated with the phylogenetic data is opening opportunities to resolve the evolutionary and ecological circumstances that have led to species diversification and speciation. This approach is opening new prospects for plant systematics that go beyond the exploratory interpretation of phylogenetic trees providing hypotheses into the “how” and “why” of biodiversity. However, identified mechanisms usually require to be verified with follow-up studies. Despite that systematics is fundamental to biodiversity, funding of studies is still a challenge despite ever-increasing future challenges^2^. However, with the ongoing digitization efforts such as collectomics^86^, integration of different kinds of already existing data with comparably affordable ‘omics such as metabolomics needs to be engaged. Integration can help to overcome these limitations while also delivering new insights. The methodology presented herein is not limited to complex-thallose liverworts and can principally be applied to any kind of organism.

## Methods

The methods are presented in detail in our related work^32,87^ and describe the steps in producing the data, including full descriptions of the experimental design, data acquisition, data science procedures, computational processing, data analysis and biostatistics. In short, samples of 16 specimens of complex-thallose liverworts were collected at field locations in Southern Sweden in September 2022 and in Germany in October 2022. Bioimaging was performed using macro- and microscopy in the lab. Plant material was isolated, washed and shock-frozen (for untargeted metabolomics analyses using LC-MS-MS), or dried (DNA marker sequence analyses). Voucher specimens were kept in the herbarium Haussknecht Jena (barcodes: JE04010739-JE04010754).

### Construction of phylogenetic trees

The phylogenetic reconstructions for the plastid region of *trn*LF were conducted with RAxML-NG version 1.2.051 using the model GTR with discrete GAMMA (GTR+G). Alpha of phylogenetic trees was estimated using Maximum Likelihood (100) and clade support were estimated using bootstrapping (1000). Additional nucleotide sequences of the investigated species were obtained from Genbank^88^, if available, using the R package rentrez version 1.2.4 and included in the construction of the phylogenetic tree. States of morphometric and chemometric characters were discretized into 8 states using the gap weighting algorithm^89,90^. Phylogenetic reconstructions of the resulting data were accomplished using RAxML-NG, but with using a multistate model (MULTI8_GTR). Dendrograms were constructed in R using the plotBS function of the R package phangorn version 2.12.1. Co-phylogenetic plots were constructed using the cophylo function from the phytools version 2.4-4 package. Consensus trees were constructed using the MAST algorithm in IQTREE version 2.2.2.6 using the “GTR+G+T” discrete gamma substitution model and with estimating branch lengths post-hoc on the consensus tree using the original alignment. Plots were accomplished with the packages ggtree version 3.16.3, ggplot2 version 4.0.1 and cowplot version 1.2.0.

### Explorative multivariate analyses

For explorative analyses, principal components analysis (PCA) was performed using the prcomp function in R. Plots were accomplished using the autoplot function of the ggplot2 version 4.0.1 and ggfortify version 0.4.19 packages. In order to assess the influence of different study factors, variation partitioning was performed using the function varpart in the package vegan 2.7-1.

### Data mining

To evaluate how much variation evolutionary and ecological factors explain in the data, the function fitExtractVarPartModel of the R package variancePartition version 1.38.0 was used. Violin plots were accomplished using the function plotVarPart of the same R package. To evaluate the significance of morpho- and chemometric characters, variable selection was employed with Random Forest (RF) using the randomForest version 4.7-1 and caret version 7.0-1 packages. A prediction model was trained using the train function from the caret package, and variable importance values were extracted from the model using the varImp function from the R package caret. Variables were selected (hence, were considered significant) when their variable importance was above 0.95. In order to visualize significant relationships of the selected variables, heatmaps were generated from the selected variables using the gplots package version 3.1.3.1. To evaluate the performance of the fitted models, 10-fold cross-validation (package mltest version 1.0.1), and the Receiver Operating Characteristic (ROC) and PR (Precision and Recall) curves using the functions plot.roc and ci.se from the pROC 1.18.5 package and the function pr.curve from the PRROC 1.3.1 package were utilized^91^.

## Data availability

- Bioimaging data: The raw images (Canon CR3-format), the pre-processed images (16-bit TIFF-format) and the contextual metadata were deposited to BioStudies under the identifier S-BIAD824 (https://www.ebi.ac.uk/biostudies/studies/S-BIAD824). The data record consists of a total of 102’375 individual raw image files partitioned into 16 samples. The entire data record has a total size of approx. 8 TB. The pre-processed and processed images along with metadata were deposited to the Image Data Resource (IDR) repository under the identifier idr0157 (https://idr.openmicroscopy.org/search/?query=Name:157). The data record consists of a total of 1127 pre-processed and 157 fully processed imaged files. The data record has a total size of approx. 12 TB. Fully segmented images showing the species are available in Zenodo (https://doi.org/10.5281/zenodo.10683968).
- Chemometric data: Raw metabolite profiles (zipped vendor data and in mzML format) have been made available in MetaboLights under the study identifier MTBLS2239 (https://www.ebi.ac.uk/metabolights/editor/MTBLS2239). The dataset includes 48 metabolite profiles in positive and negative modes, QC and blank profiles, metabolite feature tables (MAF) and metadata.
- Extracted tables and code used for analysis were deposited to Zenodo (https://doi.org/10.5281/zenodo.21478658). The dataset includes the RData containing all data objects and the R vignette containing code to reproduce to plots used in this study.
- Sequencing data were deposited to the European Nucleotide Archive (ENA) and are available under the study identifier ERP155252 (accession PRJEB70317) (https://www.ebi.ac.uk/ena/browser/view/PRJEB70317). Raw reads are available under the sample identifiers SAMEA114863468-SAMEA114863483.
- Metadata to voucher specimens are available at the virtual herbarium JACQ with the following identifiers: JE4010742, JE4010741, JE4010739, JE4010740, JE4010749, JE4010752, JE4010747, JE4010743, JE4010753, JE4010748, JE4010746, JE4010754, JE4010745, JE4010744, JE4010750, JE4010751.

## Acknowledgments

KP acknowledges the support of iDiv (funded by the German Research Foundation, DFG-FZT 118, 202548816).

## Author contributions

KP conceptualized and performed the entire study and wrote the manuscript.

## Notes

https://www.ebi.ac.uk/biostudies/studies/S-BIAD824

https://idr.openmicroscopy.org/search/?query=Name:157

https://doi.org/10.5281/zenodo.10683968

https://www.ebi.ac.uk/metabolights/editor/MTBLS2239

https://doi.org/10.5281/zenodo.21478658

https://www.ebi.ac.uk/ena/browser/view/PRJEB70317

### Competing Interest Statement

The authors have declared no competing interest.

## References

1. Schlick-Steiner, B. C. et al. Integrative Taxonomy: A Multisource Approach to Exploring Biodiversity. Annu. Rev. Entomol. 55, 421–438 (2010).

2. Orr, M. C. et al. Taxonomy must engage with new technologies and evolve to face future challenges. *Nat*. Ecol. Evol. 5, 3–4 (2020).

3. Will, K. W., Mishler, B. D. & Wheeler, Q. D. The Perils of DNA Barcoding and the Need for Integrative Taxonomy. Syst. Biol. 54, 844–851 (2005).

4. Printzen, C. Lichen Systematics: The Role of Morphological and Molecular Data to Reconstruct Phylogenetic Relationships. in Progress in Botany, Vol. 71 (eds Lüttge, U. E., Beyschlag, W., Büdel, B. & Francis, D.) vol. 71 233–275 (Springer Berlin Heidelberg, Berlin, Heidelberg, 2010).

5. Padial, J. M. & Miralles, A. RTehvieew integrative future of taxonomy. (2010).

6. Peters, K. & König-Ries, B. Reference bioimaging to assess the phenotypic trait diversity of bryophytes within the family Scapaniaceae. Sci. Data 9, 598 (2022).

7. Peters, K., Blatt-Janmaat, K. L., Tkach, N., Van Dam, N. M. & Neumann, S. Untargeted Metabolomics for Integrative Taxonomy: Metabolomics, DNA Marker-Based Sequencing, and Phenotype Bioimaging. Plants 12, 881 (2023).

8. Peters, K., Poeschl, Y., Blatt-Janmaat, K. L. & Uthe, H. Ecometabolomics Studies of Bryophytes. in Bioactive Compounds in Bryophytes and Pteridophytes (ed. Murthy, H. N.) 1–43 (Springer International Publishing, Cham, 2022). doi:10.1007/978-3-030-97415-2_30-1.

9. Renner, M. A. Opportunities and challenges presented by cryptic bryophyte species. Telopea 23, 41–60 (2020).

10. Flores, J. R., Suárez, G. M. & Hyvönen, J. Reassessing the role of morphology in bryophyte phylogenetics: Combined data improves phylogenetic inference despite character conflict. Mol. Phylogenet. Evol. 143, 106662 (2020).

11. Ludwiczuk, A., Sukkharak, P., Gradstein, R., Asakawa, Y. & Glowniak, K. Chemical relationships between liverworts of the family Lejeuneaceae (Porellales, Jungermanniopsida). Nat. Prod. Commun. 8, 1515–1518 (2013).

12. Gradstein, S., Matsuda, R. & Asakawa, Y. A chemotaxonomic survey of terpenoids and aromatic compounds in the Lejeuneaceae (Hepaticae). Contrib. Monogr. Lejeuneaceae Subfamily Ptychanthoideae 80, 63–86 (1985).

13. Figueiredo, A. C. et al. Liverwort *Radula* species from Portugal: chemotaxonomical evaluation of volatiles composition. Flavour Fragr. J. 24, 316–325 (2009).

14. Hegnauer, R. *Chemotaxonomie der Pflanzen*. (Birkhäuser Basel, Basel, 1992). doi:10.1007/978-3-0348-8649-9.

15. Stravs, M. A. A Decade of Computational Mass Spectrometry from Reference Spectra to Deep Learning. CHIMIA 78, 525–530 (2024).

16. Bechteler, J. et al. Comprehensive phylogenomic time tree of bryophytes reveals deep relationships and uncovers gene incongruences in the last 500 million years of diversification. Am. J. Bot. ajb2.16249 (2023) doi:10.1002/ajb2.16249.

17. Bowman, J. L. et al. Insights into Land Plant Evolution Garnered from the Marchantia polymorpha Genome. Cell 171, 287–304.e15 (2017).

18. Davies, K. M. et al. The Evolution of Flavonoid Biosynthesis: A Bryophyte Perspective. Front. Plant Sci. 11, 7 (2020).

19. Wheeler, J. A. Molecular Phylogenetic Reconstructions of the Marchantioid Liverwort Radiation. The Bryologist 103, 314–333 (2000).

20. Flores, J. R. et al. Dating the evolution of the complex thalloid liverworts (Marchantiopsida): total evidence dating analysis supports a Late Silurian Early Devonian origin and post Mesozoic morphological stasis. New Phytol. 240, 2137–2150 (2023).

21. Navas Romero, A. L., Herrera Moratta, M. A., Vento, B., Rodriguez, R. A. & Martínez Carretero, E. E. Variations in the coverage of biological soil crusts along an aridity gradient in the central-west Argentina. Acta Oecologica 109, 103671 (2020).

22. Weber, B. et al. What is a biocrust? A refined, contemporary definition for a broadening research community. Biol. Rev. 97, 1768–1785 (2022).

23. Seppelt, R. D., Downing, A. J., Deane-Coe, K. K., Zhang, Y. & Zhang, J. Bryophytes Within Biological Soil Crusts. in Biological Soil Crusts: An Organizing Principle in Drylands (eds Weber, B., Büdel, B. & Belnap, J.) vol. 226 101–120 (Springer International Publishing, Cham, 2016).

24. Wink, M. & Waterman, P. G. Chemotaxonomy in Relation to Molecular Phylogeny of Plants. in Annual Plant Reviews online (ed. Roberts, J. A.) 295–335 (John Wiley & Sons, Ltd, Chichester, UK, 2018). doi:10.1002/9781119312994.apr0017.

25. Heinrichs, J., Anton, H., Gradstein, S. R. & Mues, R. Systematics ofPlagiochila sect.Glaucescentes Carl (Hepaticae) from tropical America: A morphological and chemotaxonomical approach. Plant Syst. Evol. 220, 115–138 (2000).

26. *Chemical Constituents of Bryophytes: Bio- and Chemical Diversity, Biological Activity, and Chemosystematics*. (Springer Verlag, Wien ; New York, 2013).

27. Pöltl, M. et al. Unnoticed diversity in the *Riccia glauca-bifurca* group (Ricciaceae, Marchantiales): morphological differentiation and phylogeny of *R. gothica* and *R. pusilla* in Europe. Plant Biosyst. - Int. J. Deal. Asp. Plant Biol. 159, 548–562 (2025).

28. Van Zuijlen, K. et al. Bryophytes of Europe Traits (BET) data set: A fundamental tool for ecological studies. J. Veg. Sci. 34, e13179 (2023).

29. Dührkop, K. et al. SIRIUS 4: a rapid tool for turning tandem mass spectra into metabolite structure information. Nat. Methods 16, 299–302 (2019).

30. Dührkop, K. et al. Systematic classification of unknown metabolites using high-resolution fragmentation mass spectra. Nat. Biotechnol. 39, 462–471 (2021).

31. Hoffmann, M. A. et al. High-confidence structural annotation of metabolites absent from spectral libraries. Nat. Biotechnol. 40, 411–421 (2022).

32. Peters, K., Van Dam, N. M. & Neumann, S. Essential Molecular Variables of liverworts in biological soil crusts. https://doi.org/10.64898/2026.07.16.738858 (2026) doi:10.64898/2026.07.16.738858.

33. Sack, L. & Buckley, T. N. Trait Multi-Functionality in Plant Stress Response. Integr. Comp. Biol. 60, 98–112 (2020).

34. Kim, H. W. et al. NPClassifier: A Deep Neural Network-Based Structural Classification Tool for Natural Products. J. Nat. Prod. 84, 2795–2807 (2021).

35. Wong, T. K., et al. MAST: Phylogenetic Inference with Mixtures Across Sites and Trees.

36. Hyvönen, J., Koskinen, S., Smith Merrill, G. L., Hedderson, T. A. & Stenroos, S. Phylogeny of the Polytrichales (Bryophyta) based on simultaneous analysis of molecular and morphological data. Mol. Phylogenet. Evol. 31, 915–928 (2004).

37. Olson, M. E. Combining Data from DNA Sequences and Morphology for a Phylogeny of Moringaceae (Brassicales).

38. Susko, E. Tests for Two Trees Using Likelihood Methods. Mol. Biol. Evol. 31, 1029– 1039 (2014).

39. Farris, J. S., Källersjö, M., Kluge, A. G. & Bult, C. Testing Significance of Incongruence. Cladistics 10, 315–319 (1994).

40. Cummings, M. P. PAUP * [Phylogenetic Analysis Using Parsimony (and Other Methods)]. in Dictionary of Bioinformatics and Computational Biology (eds Hancock, J. M. & Zvelebil, M. J.) (Wiley, 2004). doi:10.1002/0471650129.dob0522.

41. Templeton, A. R. Phylogenetic Inference from Restriction Endonuclease Cleavage Site Maps with Particular Reference to the Evolution of Humans and the Apes. Evolution 37, 221–244 (1983).

42. Zander, R. H. Reformulation of the Foundations of Taxonomy Using Evolutionary Mechanics. Taxonomy 6, 41 (2026).

43. Renzaglia, K. S. et al. Bryophyte phylogeny: Advancing the molecular and morphological frontiers. The Bryologist 110, 179–213 (2007).

44. Wright, A. M. A Systematist’s Guide to Estimating Bayesian Phylogenies From Morphological Data. Insect Syst. Divers. 3, 2 (2019).

45. Mark, K. et al. Lichen chemistry is concordant with multilocus gene genealogy in the genus Cetrelia (Parmeliaceae, Ascomycota). Fungal Biol. 123, 125–139 (2019).

46. Xiang, Y.-L., Shen, C., Ma, W.-Z. & Zhu, R.-L. Molecular Phylogenetics and the Evolution of Morphological Complexity in Aytoniaceae (Marchantiophyta). Plants 13, 1053 (2024).

47. Furumizu, C. & Aalen, R. B. Peptide signaling through leucine rich repeat receptor kinases: insight into land plant evolution. New Phytol. 238, 977–982 (2023).

48. During, H. J. & Van Tooren, B. F. Recent developments in bryophyte population ecology. Trends Ecol. Evol. 2, 89–93 (1987).

49. Gornall, J. L., Woodin, S. J., Jónsdóttir, I. S. & van der Wal, R. Balancing positive and negative plant interactions: how mosses structure vascular plant communities. Oecologia 166, 769–782 (2011).

50. Kimura, S. & Ariyama, H. INFLUENCE OF PANTOTHENIC ACID ANALOGUES UPON THE GERMINATION OF HIGHER PLANTS. J. Vitaminol. (Kyoto) 9, 9–16 (1963).

51. Forde, B. G. & Lea, P. J. Glutamate in plants: metabolism, regulation, and signalling. J. Exp. Bot. 58, 2339–2358 (2007).

52. Brambilla, M. et al. Glutamate dehydrogenase in Liverworld—A study in selected species to explore a key enzyme of plant primary metabolism in Marchantiophyta. Physiol. Plant. 175, e14071 (2023).

53. Rabbani, N., Al-Motawa, M. & Thornalley, P. J. Protein Glycation in Plants—An Under-Researched Field with Much Still to Discover. Int. J. Mol. Sci. 21, 3942 (2020).

54. Thimmappa, R., Geisler, K., Louveau, T., O’Maille, P. & Osbourn, A. Triterpene Biosynthesis in Plants. Annu. Rev. Plant Biol. 65, 225–257 (2014).

55. Nandy, S. & Dey, A. Bibenzyls and bisbybenzyls of bryophytic origin as promising source of novel therapeutics: pharmacology, synthesis and structure-activity. DARU J. Pharm. Sci. https://doi.org/10.1007/s40199-020-00341-0 (2020) doi:10.1007/s40199-020-00341-0.

56. Commisso, M. et al. Bryo-Activities: A Review on How Bryophytes Are Contributing to the Arsenal of Natural Bioactive Compounds against Fungi. Plants 10, 203 (2021).

57. Gerson, E. A., Kelsey, R. G. & St Clair, J. B. Genetic variation of piperidine alkaloids in Pinus ponderosa: a common garden study. Ann. Bot. 103, 447–457 (2009).

58. Sobieszczuk-Nowicka, E., Arasimowicz-Jelonek, M., Tanwar, U. K. & Floryszak-Wieczorek, J. Plant homocysteine, a methionine precursor and plant’s hallmark of metabolic disorders. Front. Plant Sci. 13, 1044944 (2022).

59. Wang, C. et al. Revisiting the SAR of the Antischistosomal Aryl Hydantoin (Ro 13-3978). J. Med. Chem. 59, 10705–10718 (2016).

60. Yang, B., Liu, H., Yang, J., Gupta, V. K. & Jiang, Y. New insights on bioactivities and biosynthesis of flavonoid glycosides. Trends Food Sci. Technol. 79, 116–124 (2018).

61. Behr, M., Neutelings, G., El Jaziri, M. & Baucher, M. You Want it Sweeter: How Glycosylation Affects Plant Response to Oxidative Stress. Front. Plant Sci. 11, 571399 (2020).

62. Zhu, T.-T. et al. Functional characterization of UDP-glycosyltransferases from the liverwort Plagiochasma appendiculatum and their potential for biosynthesizing flavonoid 7-O-glucosides. Plant Sci. 299, 110577 (2020).

63. Zhu, T. et al. Functional specialization of two UDP glycosyltransferases MpUGT735A2 and MpUGT743A1 in the liverworts *Marchantia polymorpha*. J. Cell. Physiol. 238, 2499– 2511 (2023).

64. Zinsmeister, H. D., Becker, H. & Eicher, T. Bryophytes, a Source of Biologically Active, Naturally Occurring Material? Angew. Chem. Int. Ed. Engl. 30, 130–147 (1991).

65. Kulshrestha, S. et al. Stress, senescence, and specialized metabolites in bryophytes. J. Exp. Bot. 73, 4396–4411 (2022).

66. Berland, H. et al. Auronidins are a previously unreported class of flavonoid pigments that challenges when anthocyanin biosynthesis evolved in plants. Proc. Natl. Acad. Sci. 116, 20232–20239 (2019).

67. Zhou, Y. et al. Auronidin flavonoid pigments are a central component of the response of Marchantia polymorpha to carbon/nitrogen imbalance. Environ. Exp. Bot. 105862 (2024) doi:10.1016/j.envexpbot.2024.105862.

68. Markham, K. R. & J. Porter, L. Evidence of biosynthetic simplicity in the flavonoid chemistry of the ricciaceae. Phytochemistry 14, 199–201 (1975).

69. Kohn, G., Vandekerkhove, O., Hartmann, E. & Beutelmann, P. Acetylenic fatty acids in the ricciaceae (hepaticae). Phytochemistry 27, 1049–1051 (1988).

70. Oren, A. & Gunde-Cimerman, N. Mycosporines and mycosporine-like amino acids: UV protectants or multipurpose secondary metabolites? FEMS Microbiol. Lett. 269, 1–10 (2007).

71. Zulfiqar, M. et al. Untargeted metabolomics to expand the chemical space of the marine diatom Skeletonema marinoi. Front. Microbiol. 14, 1295994 (2023).

72. Kumar, M., Kuzhiumparambil, U., Pernice, M., Jiang, Z. & Ralph, P. J. Metabolomics: an emerging frontier of systems biology in marine macrophytes. Algal Res. 16, 76–92 (2016).

73. Barbehenn, R. V. & Peter Constabel, C. Tannins in plant–herbivore interactions. Phytochemistry 72, 1551–1565 (2011).

74. Niklas, K. J., Cobb, E. D. & Matas, A. J. The evolution of hydrophobic cell wall biopolymers: from algae to angiosperms. J. Exp. Bot. 68, 5261–5269 (2017).

75. Markham, K. R., Porter, L. J., Mues, R., Zinsmeister, H. D. & Brehm, B. G. Flavonoid variation in the liverwort Conocephalum conicum: Evidence for geographic races. Phytochemistry 15, 147–150 (1976).

76. Nascimento, L. B. D. S. & Tattini, M. Beyond Photoprotection: The Multifarious Roles of Flavonoids in Plant Terrestrialization. Int. J. Mol. Sci. 23, 5284 (2022).

77. Zanzoni, A., Ribeiro, D. M. & Brun, C. Understanding protein multifunctionality: from short linear motifs to cellular functions. Cell. Mol. Life Sci. 76, 4407–4412 (2019).

78. Horn, A. et al. Natural Products from Bryophytes: From Basic Biology to Biotechnological Applications. Crit. Rev. Plant Sci. 40, 191–217 (2021).

79. Plaszkó, T. et al. Correlations Between the Metabolome and the Endophytic Fungal Metagenome Suggests Importance of Various Metabolite Classes in Community Assembly in Horseradish (Armoracia rusticana, Brassicaceae) Roots. Front. Plant Sci. 13, 921008 (2022).

80. Dixon, R. A. & Paiva, N. L. Stress-Induced Phenylpropanoid Metabolism. Plant Cell 1085–1097 (1995) doi:10.1105/tpc.7.7.1085.

81. Hui, R. et al. Effects of enhanced ultraviolet-B radiation, water deficit, and their combination on UV-absorbing compounds and osmotic adjustment substances in two different moss species. Environ. Sci. Pollut. Res. 25, 14953–14963 (2018).

82. De Vries, S. et al. The evolution of the phenylpropanoid pathway entailed pronounced radiations and divergences of enzyme families. Plant J. 107, 975–1002 (2021).

83. Diaz, S., Cabido, M. & Casanoves, F. Plant functional traits and environmental filters at a regional scale. J. Veg. Sci. 9, 113–122 (1998).

84. Li, F.-W. et al. Anthoceros genomes illuminate the origin of land plants and the unique biology of hornworts. Nat. Plants 6, 259–272 (2020).

85. Bolger, M. E., Arsova, B. & Usadel, B. Plant genome and transcriptome annotations: from misconceptions to simple solutions. Brief. Bioinform. bbw135 (2017) doi:10.1093/bib/bbw135.

86. Sigwart, J. D. et al. Collectomics – towards a new framework to integrate museum collections to address global challenges. Nat. Hist. Collect. Museomics 2, 1–20 (2025).

87. Peters, K., Ziegler, J. & Neumann, S. Estimating Essential Phenotypic and Molecular Traits from Integrative Biodiversity Data. http://biorxiv.org/lookup/doi/10.1101/2024.04.02.587699 (2024) doi:10.1101/2024.04.02.587699.

88. Benson, D. A. et al. GenBank. Nucleic Acids Res. 41, D36–D42 (2012).

89. Thiele, K. The Holy Grail of the Perfect Character: the Cladistic Treatment of Morphometric Data. Cladistics 9, 275–304 (1993).

90. Garcia-Cruz, J. & Sosa, V. Coding Quantitative Character Data for Phylogenetic Analysis: A Comparison of Five Methods. Syst. Bot. 31, 302–309 (2006).

91. Grau, J., Grosse, I. & Keilwagen, J. PRROC: computing and visualizing precision-recall and receiver operating characteristic curves in R. Bioinformatics 31, 2595–2597 (2015).

